## Supplementary Materials for "Rhythmic interactions between the mediodorsal thalamus and prefrontal cortex precede human visual perception"

**This PDF file includes:**

Figs. S1 to S12
Table S1

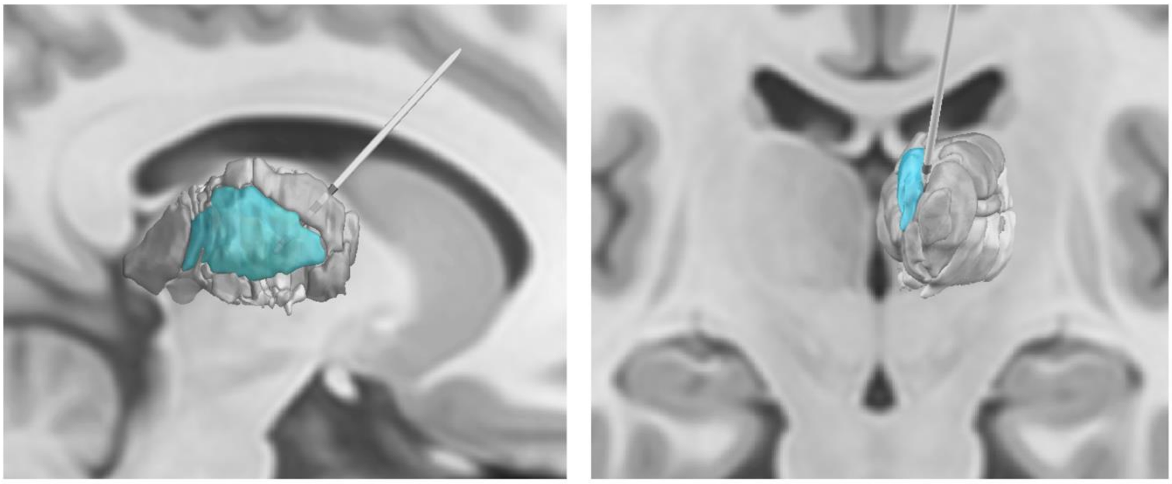

**Supplementary Fig. 1. Thalamic implantation. Unilateral depiction of the** mediodorsal thalamus (aqua) in the context of other thalamic nuclei (rendered in grey).

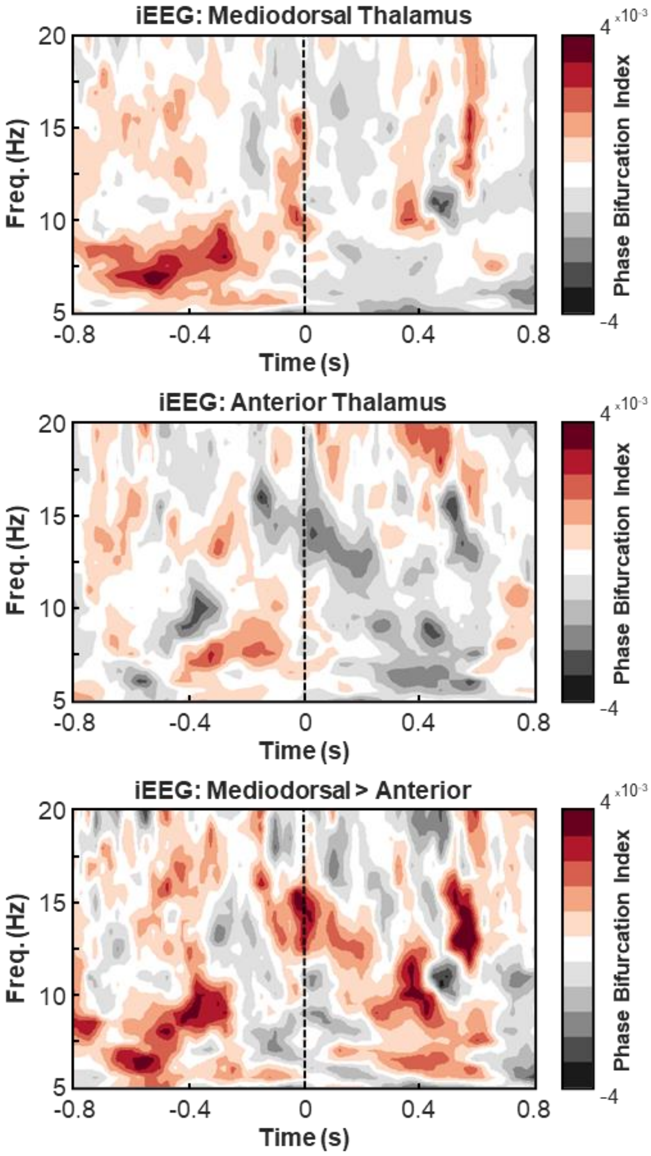


**Supplementary Fig. 2. Comparison of phase bifurcation in the mediodorsal and anterior thalamic nuclei.** Top: Time-frequency representation depicting mean mediodorsal thalamic phase bifurcation across patients (as measured with iEEG). Higher values indicate greater phase bifurcation. Time at 0s represents onset of the target. Duplicated from Fig. 1C. Middle: Time-frequency representation depicting mean anterior thalamic phase bifurcation across patients (as measured with iEEG). While, descriptively speaking, there are increases in the phase bifurcation index in the low frequencies, this was not statistically significant ($p > 0.5$). Bottom: Time-frequency representation depicting the difference in mean mediodorsal thalamic phase bifurcation and mean anterior thalamic phase bifurcation across patients (as measured with iEEG).

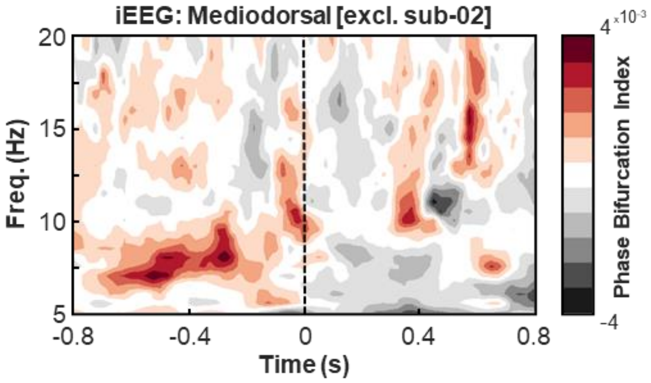


**Supplementary Fig. 3. Phase bifurcation in the mediodorsal thalamus after excluding participant 2.** As shown in figure 1e, participant 2 produced a larger phase bifurcation index at a lower peak frequency than others. To see if this drives the effect presented in figure 1c, we re-plotted this figure after excluding participant 2. This plot demonstrates the same narrowband, low-frequency, pre-stimulus increase in phase bifurcation seen in figure 1c.

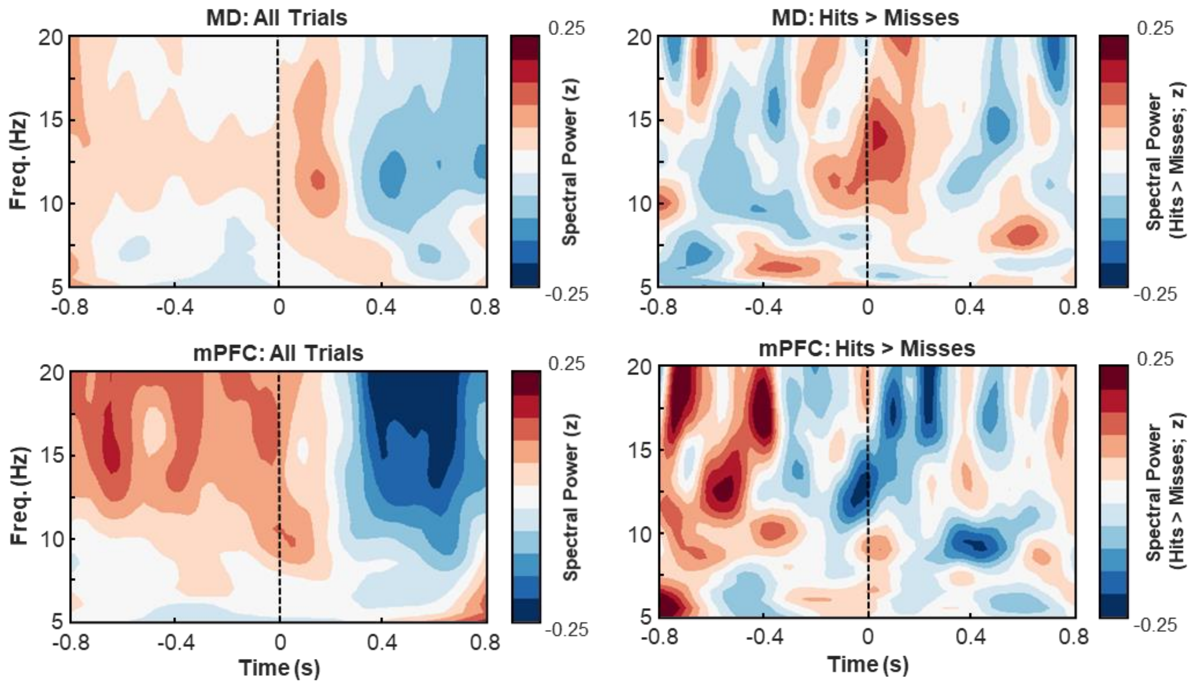


**Supplementary Fig. 4. Spectral power changes in the mediodorsal thalamus and medial prefrontal cortex.** Spectral power within the mediodorsal thalamus did not change as a function of perceptual performance (top right; $t(5) = 2.92$, $p_{clus} = 0.453$, $BF_{10} = 2.83$). Similarly, spectral power within the medial prefrontal cortex did not change as a function of perceptual performance (bottom right; $t(5) = -3.87$, $p_{clus} = 0.250$, $BF_{10} = 6.11$). Spectral power was computed using the same time-frequency decomposition parameters as those used for the phase bifurcation analyses, and the resulting power spectra for hits and misses were directly contrasted in a cluster-based, permutation dependent-samples t-test.

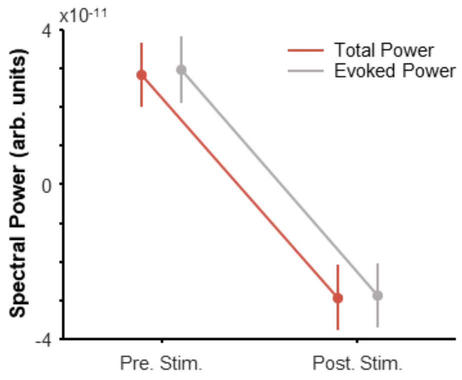


**Supplementary Fig. 5. Phase reset analysis.** To test whether the phase of ongoing activity reset following stimulus onset, we computed low-frequency spectral power (6 to 9Hz; in steps of 1Hz) just before stimulus onset (-400 to 0ms; in steps of 25ms) and just after stimulus onset (0 to 400ms; in steps of 25ms) using 6-cycle wavelets. We conducted this spectral decomposition twice: first, on single trials before averaging the result across trials (i.e., total power), and second, on the trial-averaged amplitude (i.e., evoked power). If phase resets after stimulus onset, then phase should align across trials after stimulus onset, which will present as an increase in total power for post-stimulus activity relative to pre-stimulus activity. In contrast, no change in spectral power will be observed on the single trial level (for further details, see 20). To statistically appraise the effect, we conducted a 2x2 repeated measures ANOVA to probe how spectral power changed as a function of epoch (pre- vs. post-stimulus) and decomposition method (single trial decomposition vs. trial-averaged decomposition). No significant interaction was observed ($F(1, 5) = 1.04$, $p = 0.355$), suggesting that phase did not reorganise consistently across participants.

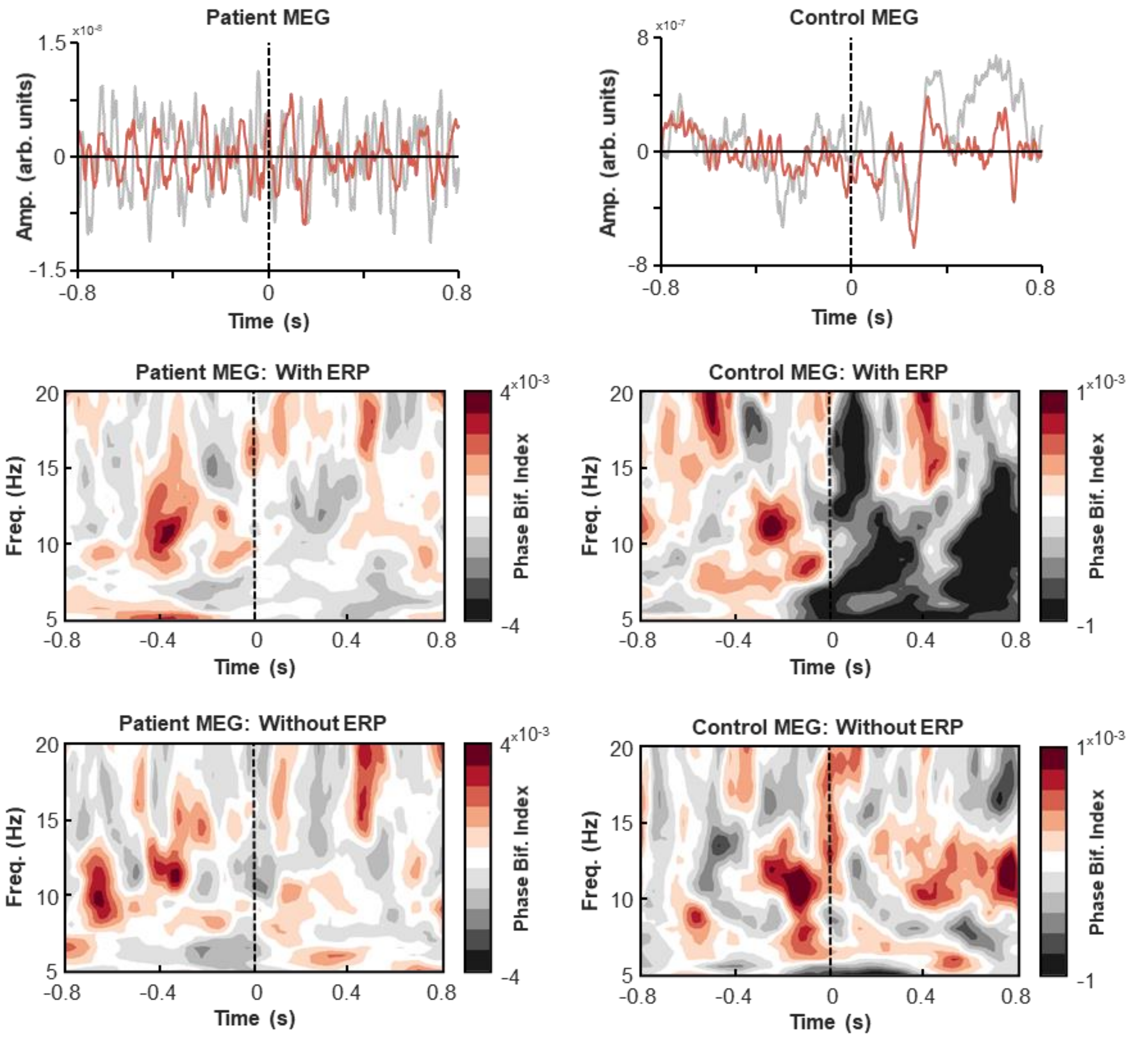


**Supplementary Fig. 6.** Influence of event-related potential on phase-bifurcation measures. Differences in the post-stimulus phase bifurcation index (PBI) can be observed between the patient and control MEG recordings, with the latter showing a temporally and spectrally broad negative effect following stimulus onset. PBI is sensitive to condition-specific differences in evoked responses, which present as large negative PBI values (Busch et al., 2009), leading to the speculation that the decrease in the control sample reflects a condition-specific difference in the evoked response. To formally test these ideas, we first tested for condition-specific differences in the evoked response for the two samples using cluster-based permutation tests on the post-stimulus data (0ms to +800ms). While the patient MEG showed no effect ($p = 0.481$; top left panel), a significant difference was observed for the control MEG ($p = 0.043$; top right panel). [Note that the differences in sub-5Hz activity between the samples is driven by the use of heavier filtering in the patient data to attenuate artifacts introduced by the intracranial wires]. The differences in ERPs would explain why the control PBI shows a negative post-stimulus PBI (middle right), while the patients do not (middle left). Indeed, if the trial-average ERP is subtracted from the single-trial data, the negative PBI in the control MEG data disappears (bottom right), while the patient MEG data remains, more-or-less, unchanged (bottom left).

Note that no negative PBI is observed in the mediodorsal thalamus because there was no condition-specific difference in the thalamic evoked response ($p = 0.509$).

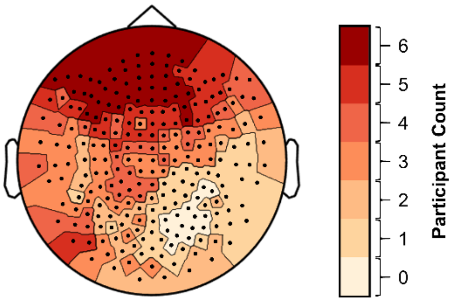


**Supplementary Fig. 7.** Topography depicting the sensors that survived artifact rejection as a function of the number of patients. The darkest red indicates that all patients retained these sensors after artifact rejection, lighter colours indicate that fewer patients retain these sensors after artifact rejection. Sensor loss arose principally over posterior regions.

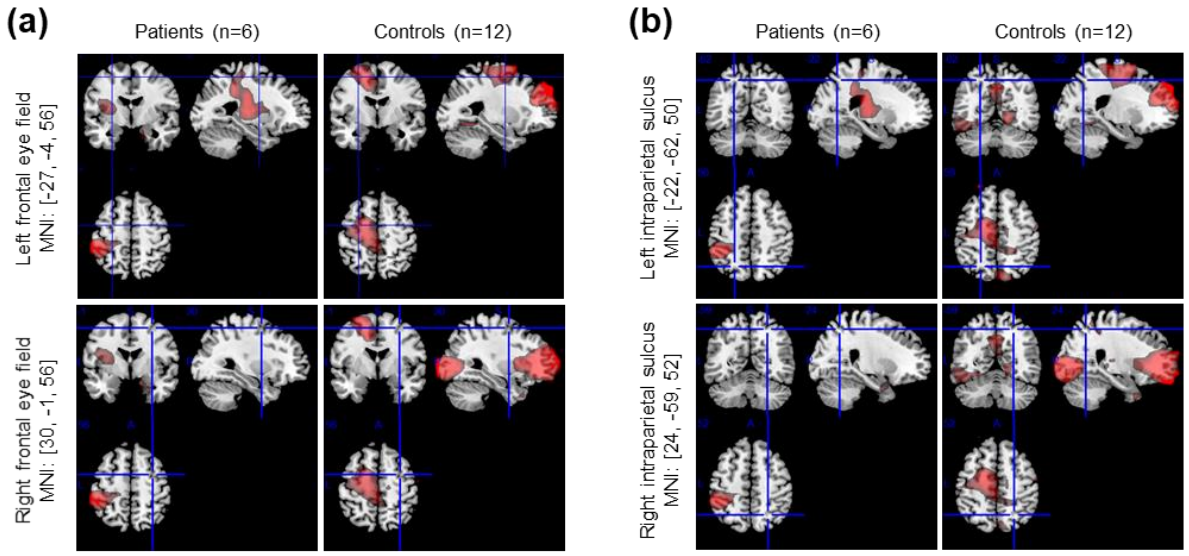


**Supplementary Fig. 8.** Previous work has demonstrated that the dorsal attention network shows similar phase bifurcation effects as the prefrontal cortex. Here, we explored whether similar effects can be observed in our data, taking two regions that have previously been shown to demonstrate rhythmic sampling: the frontal eye fields (panel a) and the intraparietal sulcus (panel b) [Fiebelkorn, Minsk & Kastner, 2019]. Panel A has been centred on MNI co-ordinates representative of the left and right frontal eye fields (Murd et al., 2020). Panel B has been centred on MNI co-ordinates representative of the intraparietal sulcus (Bray et al., 2013). While we did not observe any substantial phase bifurcation in either area for the patient data, substantial phase bifurcation was observed in the left frontal eye fields for the healthy controls. The absence of an effect in the patient data may be attributable to the fact that the externalised intracranial wires ran over the back of the head, producing large artifacts that reduce the signal-to-noise of data sampled from these regions. Consequently, the absence of an effect in the dorsal attention network of the patients should not be interpreted as an absence of an effect in the population, but simply an idiosyncratic feature of MEG recordings from patients with externalised DBS wires.

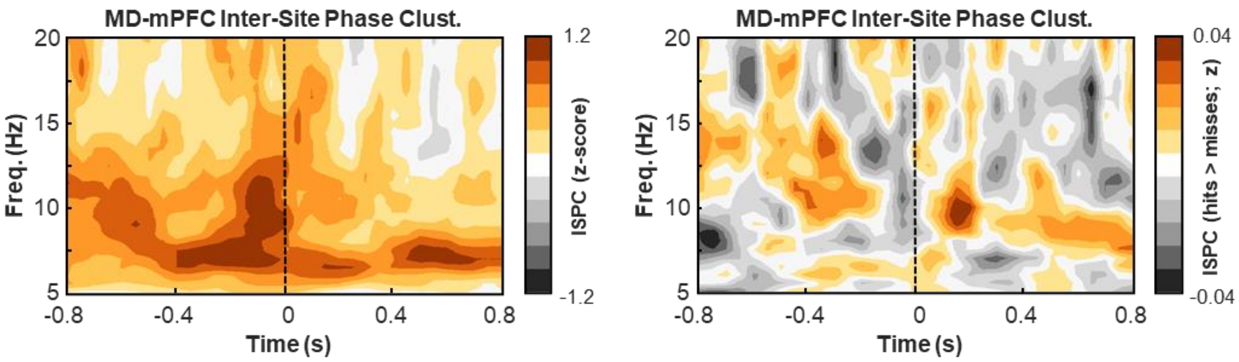


**Supplementary Fig. 9.** Inter-site phase clustering (ISPC) between the mediodorsal thalamus (MD) and medial prefrontal cortex (mPFC) across all trials (left; duplicate of Fig. 2A) and for hits relative to misses (right). No significant difference in ISPC was observed for hits relative to misses.

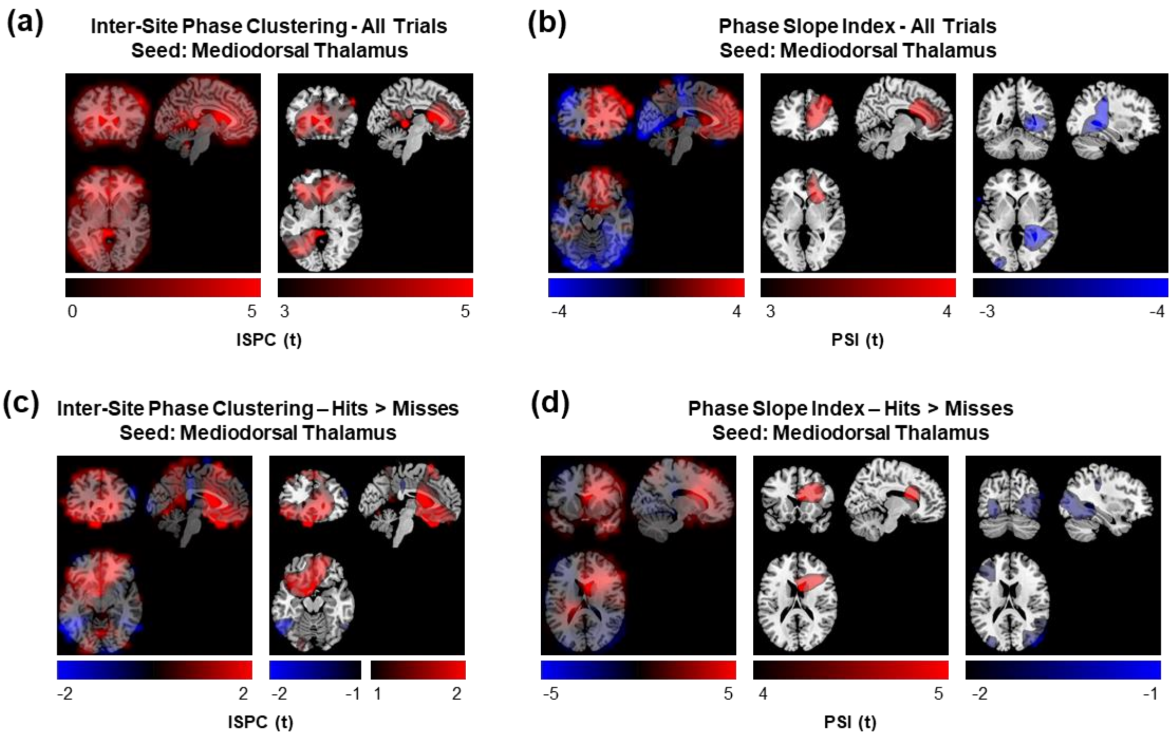


**Supplementary Fig. 10. Low-frequency thalamic connectivity across the brain.** In the main text, we had focused our iEEG-MEG connectivity analyses on the prefrontal cortex due to difficulties recording posterior sources (see supplementary figure 7). Here, however, we plot the connectivity plots for pre-stimulus low-frequency activity (-800ms to 0ms; 6-14Hz) in full. Given the limited and varying coverage of posterior sources, these effects should be interpreted with caution. **(a)** Inter-site phase clustering across all trials (left: unthresholded; right: thresholded). As reported in the main text, we observed substantial undirected connectivity between the mediodorsal thalamus and medial prefrontal cortex (mean cluster $t(5) = 19.83$, $p_{clus} < 0.001$, $BF_{10} = 2{,}218.64$). In addition, we saw a second, smaller cluster of connectivity between the ipsilateral occipital lobe and mediodorsal thalamus. When restricting the analysis space to posterior sources [and hence excluding the prefrontal cortex from analysis], this cluster was found to be significant (mean cluster $t(5) = 13.88$, $p_{clus} < 0.001$, $BF_{10} = 551.72$). **(b)** Phase slope index across all trials (left: unthresholded; middle: positive thresholding; right; negative thresholding). A positive phase slope index indicates that the cortex leads the thalamus, while a negative phase slope index indicates that the thalamus leads the cortex. As reported in the main text, we observed significant directed connectivity from the medial prefrontal cortex to the mediodorsal thalamus (mean cluster $t(5) = 5.33$, $p_{clus} < 0.001$, $BF_{10} = 16.73$). In addition, we observed directed connectivity from the mediodorsal thalamus to the occipital/parietal lobes that trended towards significance (mean cluster $t(5) = -8.15$, $p_{clus} = 0.063$, $BF_{10} = 73.96$). **(c)** Inter-site phase clustering for hits relative to misses (left: unthresholded; right: thresholded). While the difference in connectivity between conditions was greatest between the mediodorsal thalamus and medial prefrontal cortex, this was not significant (mean cluster $t(5) = 5.37$, $p_{clus} = 0.188$, $BF_{10} = 17.13$). **(d)** Phase slope index for hits relative to misses (left: unthresholded; middle: positive thresholding; right; negative thresholding). As reported in the main text, we found that directed connectivity from the medial prefrontal cortex to the mediodorsal thalamus was significantly greater for hits relative to misses (mean cluster $t(5) = 8.26$, $p_{clus} < 0.001$, $BF_{10} = 11.71$). Directed connectivity from the mediodorsal thalamus to the occipital lobe did not differ for hits relative to misses (mean cluster $t(5) = -2.80$, $p_{clus} = 0.844$, $BF_{10} = 2.54$).

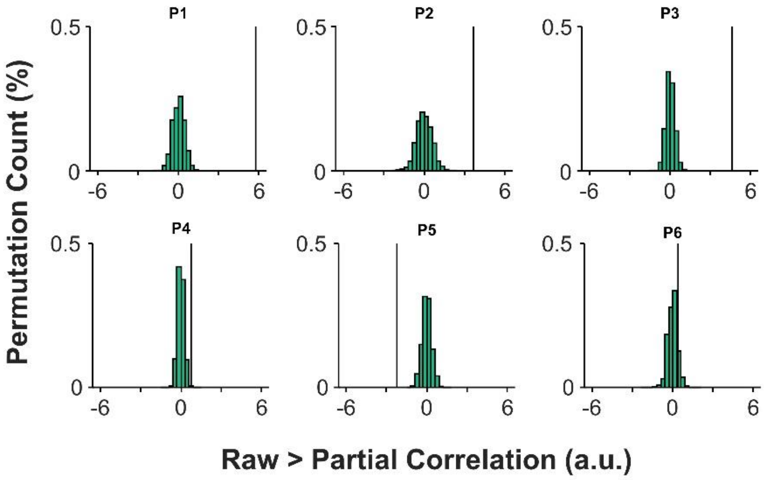


**Supplementary Fig. 11.** The difference in correlation co-efficient between medial prefrontal cortical optimal phase and perceptual performance when accounting for mediodorsal thalamic activity (chance difference represented by histogram bars; 1,000 permutations). To supplement the mediation analysis presented in the main text, we computed the correlation between medial prefrontal phase bifurcation and perceptual performance, and the partial correlation of these two variables after accounting for mediodorsal thalamic phase bifurcation. The resulting correlation coefficients were directly contrasted (that is, the raw correlation minus the partial correlation coefficient). A resulting positive value indicates that the raw coefficient was larger than the partial coefficient, which indicates that the mediodorsal thalamus explains some of the correlation between medial prefrontal phase bifurcation and perceptual performance.

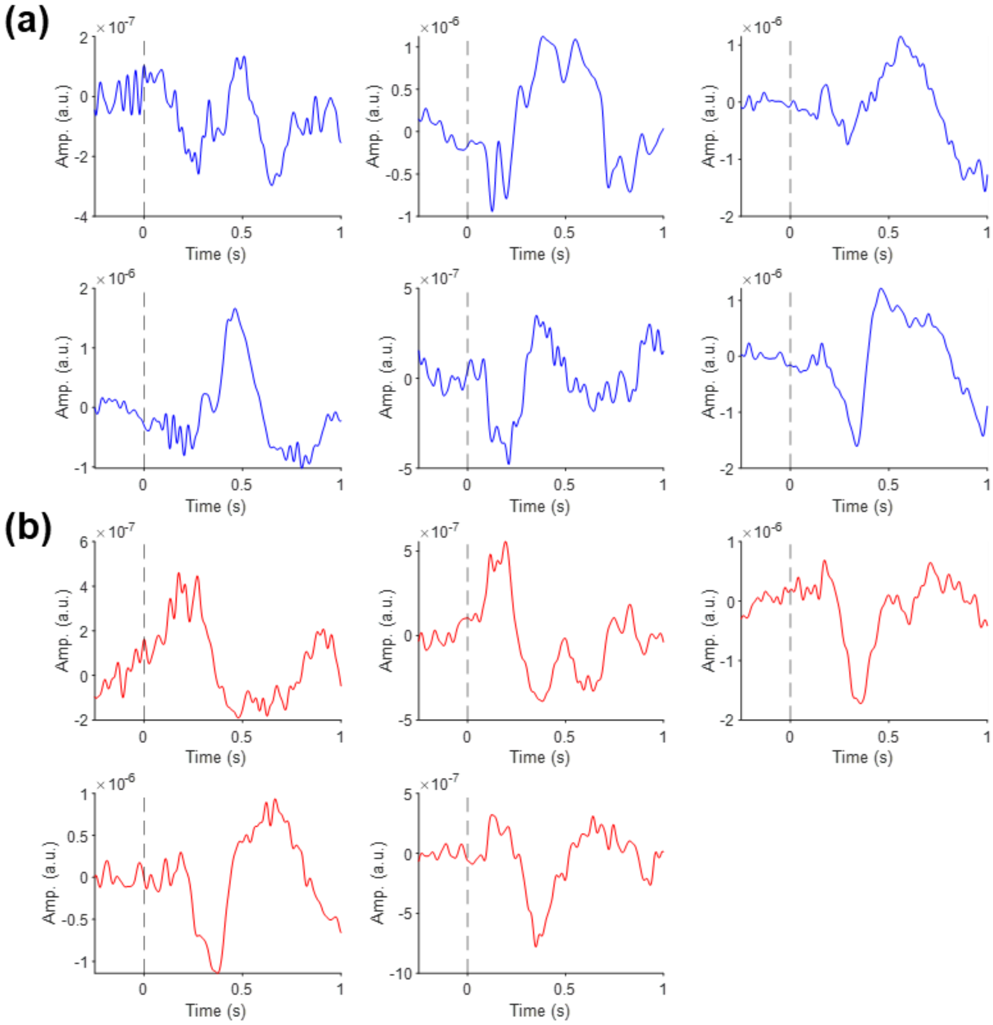


**Supplementary Fig. 12. Mean evoked response to stimulus onset across all trials. (a)** Mean mediodorsal evoked response (blue) to stimulus onset (time = 0s) for each of the six patients. **(b)** Mean anterior evoked response (red) to stimulus onset (time = 0s) for five patients (one participant had no bipolar-referenced contact within the anterior thalamus).

**Supplementary Table 1.** Location and bipolar referencing montage of intracranial electrodes. Labels were taken from the X atlas. We defined the mediodorsal thalamus (MD) as those electrodes sitting within either the *Nucleus Medialis* or *Lamella Medialis Thalami*, and the anterior thalamus (ANT) as those electrodes sitting within the *Nucleus Anterior Principalis*. Due to the small size of the ANT, we were forced to re-reference the ANT recordings of one participant (pp05) to the *Lamella Medialis Thalami.*

| Participant | Hemisphere | Contact Num. | Principle Label | MD Bipolar Pair | ANT Bipolar Pair |
|---|---|---|---|---|---|
| 1 | Left | 1 | Nucleus Medialis | | |
| | | 2 | Lamella Medialis Thalami | | |
| | | 3 | Nucleus Anterior Principalis | | |
| | | 4 | Nucleus Anterior Principalis | | |
| | Right | 1 | Nucleus Medialis | * | |
| | | 2 | Nucleus Medialis | * | |
| | | 3 | Nucleus Anterior Principalis | | * |
| | | 4 | Nucleus Anterior Principalis | | * |
| 2 | Left | 1 | Nucleus Medialis | * | |
| | | 2 | Nucleus Medialis | * | |
| | | 3 | Nucleus Anterior Principalis | | * |
| | | 4 | Nucleus Anterior Principalis | | * |
| | Right | 1 | Nucleus Medialis | | |
| | | 2 | Lamella Medialis Thalami | | |
| | | 3 | Nucleus Anterior Principalis | | |
| | | 4 | Nucleus Anterior Principalis | | |
| 3 | Left | 1 | Lamella Medialis Thalami | | |
| | | 2 | Lamella Medialis Thalami | | |
| | | 3 | VA | | |
| | | 4 | VA | | |
| | Right | 1 | Nucleus Medialis | * | |
| | | 2 | Lamella Medialis Thalami | * | |
| | | 3 | Nucleus Anterior Principalis | | * |
| | | 4 | Nucleus Anterior Principalis | | * |
| 4 | Left | 1 | Nucleus Medialis | | |
| | | 2 | Nucleus Anterior Principalis | * | |
| | | 3 | Nucleus Anterior Principalis | * | |
| | | 4 | Corpus Callosum | | |
| | Right | 1 | Nucleus Medialis | | * |
| | | 2 | Lamella Medialis Thalami | | * |
| | | 3 | Nucleus Anterior Principalis | | |
| | | 4 | CSF | | |
| 5 | Left | 1 | Nucleus Medialis | | |
| | | 2 | Nucleus Medialis | | |
| | | 3 | Nucleus Medialis | | |
| | | 4 | Nucleus Anterior Principalis | | |
| | Right | 1 | Nucleus Medialis | * | |
| | | 2 | Nucleus Medialis | * | |
| | | 3 | Lamella Medialis Thalami | | * |
| | | 4 | Nucleus Anterior Principalis | | * |
| 6 | Left | 1 | Lamella Medialis Thalami | * | |
| | | 2 | Lamella Medialis Thalami | * | |
| | | 3 | Dorso-intermedius Internus | | |
| | | 4 | Dorso-intermedius Internus | | |
| | Right | 1 | Lamella Medialis Thalami | | |
| | | 2 | Dorso-intermedius Internus | | |
| | | 3 | Dorso-intermedius Internus | | |
| | | 4 | ? | | |
